# African swine fever virus structural protein p17 inhibits cGAS-STING signaling pathway through interacting with STING

**DOI:** 10.1101/2021.06.02.446854

**Authors:** Wanglong Zheng, Nengwen Xia, Jia Luo, Sen Jiang, Jiajia Zhang, Hui Wang, Da Ao, Yulin Xu, Xueliang Liu, Qi Shao, Qi Cao, Youwen Zhang, Nanhua Chen, Quan Zhang, Jiansen Da, Hongjun Chen, Xiaoyu Guo, Hongfei Zhu, François Meurens, Jianzhong Zhu

## Abstract

African swine fever (ASF) is highly contagious, causes high mortality in domestic and feral swine, and has a significant economic impact on the global swine industry due to the lack of a vaccine or an effective treatment. African swine fever virus (ASFV) encodes more than 150 polypeptides, which may have intricate and delicate interactions with the host for the benefit of the virus to evade the host’s defenses. However, currently, there is still a lack of information regarding the roles of the viral proteins in host cells. Here, our data demonstrated that the p17, encoded by D117L gene could suppress porcine cGAS-STING signaling pathway, exhibiting the inhibitions of TBK1 and IRF3 phosphorylations, downstream promoter activities, cellular mRNA transcriptions and ISG56 induction, and antiviral responses. Further, we found that p17 was located in endoplasmic reticulum (ER) and Golgi apparatus, and interacted with STING, perturbing it in the recruitment of TBK1 and IKKε. Additionally, it appeared that the transmembrane domain (amino acids 39–59) of p17 could be required for interacting with STING and inhibiting cGAS-STING pathway. Taken together, p17 could inhibit the cGAS-STING pathway through its interaction with STING and interference with STING in the recruitment of TBK1 and IKKε.

**Importance:** African swine fever (ASF) is a highly contagious disease in domestic and feral swine, posing significant economic impacts on the global swine industry, and the pathogen ASFV is a large icosahedral DNA virus. The innate immune cGAS-STING DNA sensing pathway plays a critical role in sensing invading ASFV and triggering antiviral responses. However, there is still a lack of information regarding the molecular mechanisms of ASFV evasion of the cGAS-STING pathway. We have analyzed the effects of whole genomic open reading frames (ORFs) of ASFV China 2018/1 on the activation of cGAS-STING pathway, and found that p17 was able to inhibit cGAS-STING mediated type I IFN production by targeting STING, altering its capacity to recruit both TBK1 and IKKε. Findings presented here will expand our knowledge on the molecular mechanisms by which ASFV counteracts the antiviral innate immunity and provide deep insights into ASF pathogenesis.

## 1. Introduction

African swine fever virus (ASFV), the causative agent of African swine fever (ASF) which is a contagious disease of domestic pigs and wild boar, has spread globally in recent years with serious economic consequences for the swine industry (1, 2). ASFV is a linear double-stranded DNA molecule that varies in length from about 170 to 193 kb and encodes for 150 to 167 open reading frames (3–5). Natural target cells of ASFV are macrophages, monocytes from lymphatic tissues, megakaryocytes, and polymorphonuclear leukocytes present in blood and bone marrow (6). In order to replicate and spread, ASFV has evolved multiple strategies to escape the host’s defense system (7). Several studies have suggested that ASFV could evade the innate immunity by modulation of a number of host cell pathways including type I IFN production and response (8), host cell gene transcription and protein synthesis (9, 10), cell proliferation and death (11, 12), and inflammatory response (13).

Accumulating evidences have demonstrated that inhibiting the production and the effects of IFN plays a critical role in ASFV pathogenesis (14). Several researches have indicated that virulent strains of ASFV, including Armenia/07, Lisboa60 and 22653/14 strains, have developed various measures to block the production and responses of IFNs in infected cells (15–17). Specifically, the virulent Armenia/07 strain was able to efficiently block the synthesis and the production of IFN-β mRNA in infected macrophages through the cGAS-STING pathway (15). The highly virulent strain Lisboa60 could inhibit IFN production in macrophages through a way independent on IRF3 modulation (16). Virulent 22653/14 strain seemed to have developed mechanisms to suppress the induction of several type I IFN genes (17). Additionally, multiple ASFV proteins, including A276R, A528R, I329L, DP96R, MGF-505-7R, and MGF360-12L, were shown to contribute to the inhibition of IFN production by targeting different pathways (14, 18–21).

P17 encoded by the ASFV D117L gene is a major structural transmembrane protein localized in the capsid and inner lipid envelope. P17 is an essential and highly abundant protein required for the assembly and maturation of the icosahedral capsid and virus viability (22, 23). Several studies have indicated that the outer capsid shell was formed by the major capsid protein (p72) and four stabilizing minor proteins (H240R, M1249L, p17, p49) (24–26). The p17 protein appears to form trimers and is located at the interface of the center gap region of three neighboring pseudo-hexameric capsomers (24). p17 closely associates with the base domain of p72, and three copies of p17 encircle each p72 trimer capsomer in the inner capsid shell, firmly anchoring p72 capsomers on the inner membrane (25). Our previous study also indicated that p17 protein could inhibit cell proliferation by inducing cell cycle arrest via ER stress-ROS pathway (23).

It is known that mammalian immune system utilizes pattern recognition receptors (PRRs) to detect pathogen-associated molecular patterns (PAMPs) from invading pathogens and activate the host innate immune response. Upon DNA virus infection of a permissive cell, viral DNA is detected by cytosolic sensors. The cytosolic cyclic GMP-AMP (cGAMP) synthase (cGAS) plays a key role in sensing cytosolic DNA and triggering the stimulator of interferon genes (STING) dependent signaling to induce type I IFNs (27). cGAS is activated upon binding with double-stranded DNA and then catalyzes the second messenger 2’3’-cGAMP generation, which in turn activates STING (28). Activated STING is translocated from endoplasmic reticulum (ER) via ER Golgi intermediate compartment (ERGIC) to the Golgi apparatus and serves as a platform for recruitment and phosphorylation of TBK1 and interferon regulatory factor 3 (IRF3) (29). TBK1 phosphorylates itself, STING and transcription factor IRF3. Phosphorylated IRF3 dimerizes and translocates to the nucleus, where it triggers the production of type I IFNs and, subsequently, the expressions of interferon-stimulated genes (ISGs), orchestrating antiviral defense mechanisms (28).

Previous studies have reported that virulent strains of ASFV could affect the production of IFN-β through the inhibition of the cGAS-STING pathway (15). However, there is still a lack of information regarding the molecular mechanisms of ASFV interferences with the cGAS-STING pathway. We have analyzed the effects of whole genomic open reading frames (ORFs) of ASFV China 2018/1 on the activation of cGAS-STING pathway, and found that p17 was able to inhibit porcine cGAS-STING mediated IFN signaling. Thus, the main aim of this study was to decipher the mechanisms underlying ASFV p17 interference with the cGAS-STING pathway.

## 2. Results

### 2.1 ASFV p17 can inhibit the DNA sensing porcine cGAS-STING signaling pathway

In order to explore the molecular mechanisms of ASFV’s immune evasion, the impact of p17 on the DNA sensing cGAS-STING pathway was analyzed. Firstly, the effects of p17 on promoters’ activations, including IFN-β, ISRE and NF-κB, were analyzed by using dual-luciferase reporter assay. The results from 293T cells showed that cGAS/STING stimulated activations of ISRE, IFN-β and NF-κB promoters were all decreased in the presence of p17 (Fig 1A and 1B). Similarly, in the p17 transfected PAMs, the polydA:dT or 2’3’-cGAMP stimulated IFN-β and ISRE promoter activations were also decreased (Fig 1C and 1D). Secondly, the effect of p17 on the cGAS-STING pathway mediated downstream cellular mRNA expressions of IFN-β, ISG15, ISG56 and IL-8 were analyzed by RT-qPCR. It appeared that p17 could inhibit polydA:dT induced downstream mRNA expressions of IFN-β, ISG15, ISG56 and IL-8 (Fig 1E).

**Figure 1.**
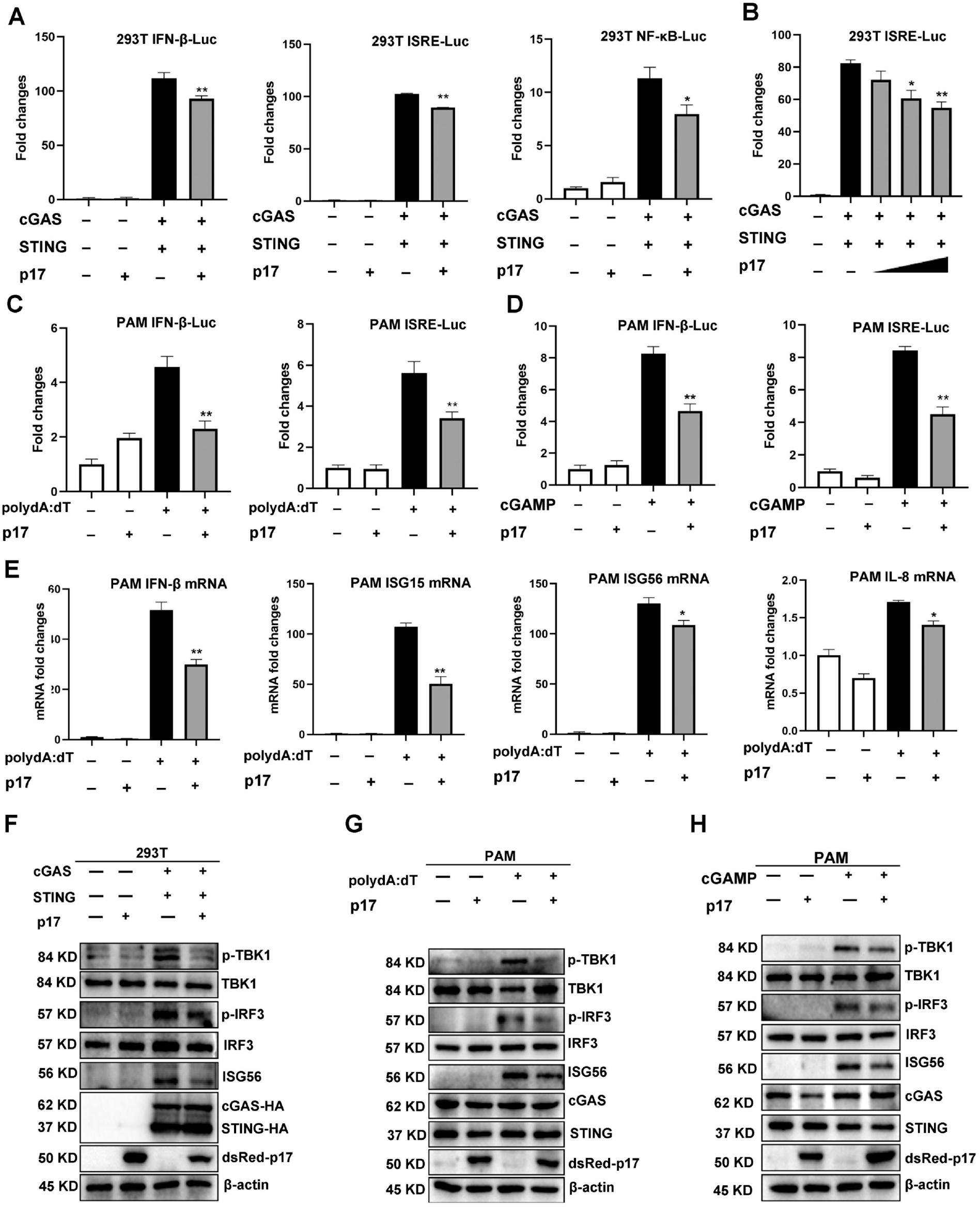
The effects of ASFV p17 on the porcine DNA sensing cGAS-STING pathway. (A and B) 293T cells grown in 96-well plate (3×10^4^ cells/well) were transfected by Lipofectamine 2000 with pcGAS (20 ng), pSTING (10 ng), p17 (10 ng), plus each Fluc reporter (10 ng) and Rluc (0.4 ng) (A), pcGAS (20 ng), pSTING (10 ng), p17 (5, 10, 20 ng), plus ISRE Fluc (10 ng) and Rluc (0.4 ng) (B), which were normalized to 50 ng/well. Twenty-four hours post-transfection, the luciferase activities were measured. (C and D) PAMs grown in 96-well plate (3×10^4^ cells/well) were transfected with p17 plasmids (20 ng), plus IFN-β-Fluc (20 ng) or ISRE-Fluc (20 ng) and Rluc (0.4 ng) for 12 h, and then stimulated by transfection of polydA:dT (1 μg/mL) (C) or 2’3’-cGAMP (2 μg/mL) (D) for 12 h, followed by the measurement of luciferase activities. (E) PAMs grown in 12-well plate (3×10^5^ cells/well) were transfected with p17 plasmid (1 μg) for 12 h and then simulated by transfection of polydA:dT (1 μg/mL) using Lipofectamine 2000 for another 12 h; the RNA expressions of IFN-β, ISG15, ISG56 and IL-8 were analyzed by RT-qPCR. (F) 293T cells grown in 12-well plate (3×10^5^ cells/well) were transfected with p17 plasmid (0.5 μg), pcGAS (0.5 μg) and pSTING (0.5 μg). Twenty-four hours post-transfection, the expressions of p-TBK1, TBK1, p-IRF3, IRF3, ISG56, cGAS and STING were analyzed by Western blotting. (G and H) PAMs grown in 12-well plate (3×10^5^cells/well) were transfected with p17 plasmid (1 μg), polydA:dT (1 μg/mL) (G) or 2’3’-cGAMP (2 μg/mL) (H) for 24 h. The signaling protein expressions were analyzed by Western blotting.

Additionally, the effects of p17 on the cGAS-STING signaling pathway were examined using Western blotting in both 293T and PAMs (Fig 1F-H). In 293T cells, the exogenous porcine cGAS-STING induced phosphorylation of TBK1 (p-TBK1) and IRF3 (p-IRF3), and downstream ISG56 induction were all inhibited by p17 (Fig 1F). In PAMs, polydA:dT and 2’3’-cGAMP both activated endogenous porcine cGAS-STING signaling, upregulating the levels of p-TBK1, p-IRF3 and downstream ISG56. With the expression of p17, the polydA:dT and 2’3’-cGAMP induced p-TBK1, p-IRF3 and downstream ISG56 were all decreased (Fig 1G and 1H). Taken together, these data clearly suggested that ASFV p17 could inhibit the DNA sensing cGAS-STING signaling pathway.

### 2.2 ASFV p17 can disturb the STING signaling mediated antiviral responses

In order to further investigate the impact of p17 on the cGAS-STING pathway, the effects of p17 on the STING signaling mediated antiviral responses were analyzed. PAMs were infected with DNA virus HSV1-GFP, and STING agonist 2’3’-cGAMP showed obvious antiviral activity. However, ASFV p17 disturbed the STING signaling mediated anti-HSV1 activity, as evidenced by fluorescence microscopy (Fig 2A), flow cytometry (Fig 2B) and Western blotting (Fig 2C). The HSV1 gB gene expression by RT-qPCR analysis and virus titration by plaque assay both showed similar results (Fig 2G and 2H).

**Figure 2.**
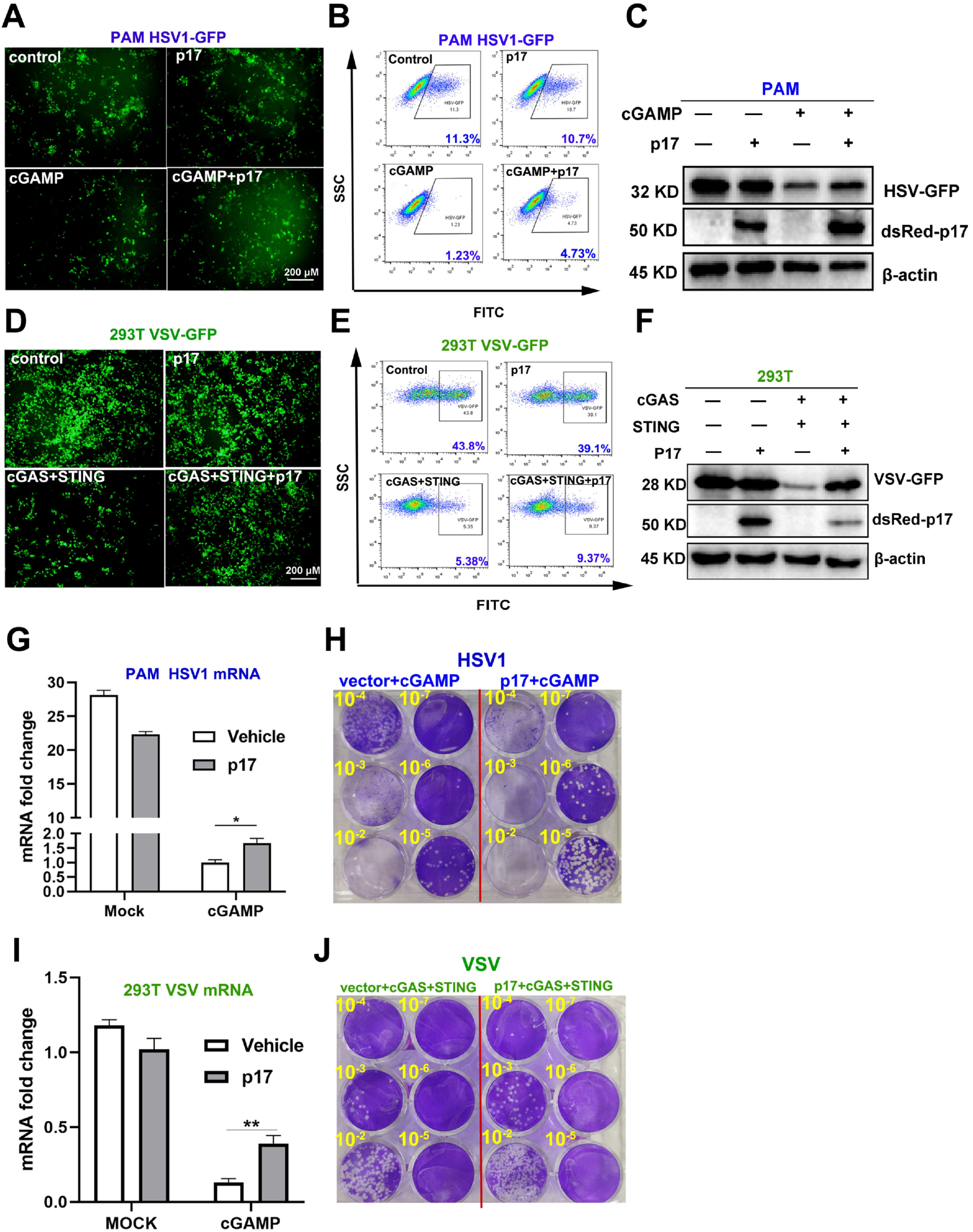
Effects of p17 on the cGAS-STING signaling mediated anti-HSV1 and anti-VSV responses. (A-C, G and H) PAMs grown in 12-well plates (3×10^5^ cells/well) were transfected with p17 plasmid (1μg) for 12 h, and next stimulated by transfection with 2’3’-cGAMP (2μg/mL) for 12 h. The cells were infected with HSV1-GFP at the MOI 0.01 for 20 h. (A) The HSV1 replicative GFP signals were observed by fluorescence microscopy. (B) The HSV1-GFP infected cells were analyzed by flow cytometry. (C) The viral GFP expressions were analyzed by Western blotting. (G) HSV1 gB gene expressions were analyzed by RT-qPCR. (H) The viral titers in the supernatants from HSV1 infected PAMs were analyzed by plaque assay. (D-F, I and J) 293T cells grown in 12-well plate (3×10^5^ cells/well) were co-transfected with p17 plasmid (0.5 μg), pcGAS (0.5 μg) and pSTING (0.5 μg) for 12 h, and then the cells were infected with VSV-GFP at the MOI = 0.001 for 8 h. The VSV replicative GFP signals were observed by fluorescence microscopy (D), analyzed by flow cytometry (E), and by Western blotting (F). The VSV glycoprotein genes were analyzed by RT-qPCR (I) and the VSV titers in supernatants from VSV infected 293T cells were measured by plaque assay (J).

The cGAS-STING signaling pathway has been shown to have broad antiviral function (30, 31). Therefore, 293T cells were infected with RNA virus VSV-GFP, and co-expressed porcine cGAS/STING triggered obvious anti-VSV activity. However, ASFV p17 disturbed the STING signaling mediated anti-VSV activity, which was evidenced by fluorescence microscopy (Fig 2D), flow cytometry (Fig 2E) and Western blotting (Fig 2F). The VSV glycoprotein gene expression by RT-qPCR analysis and virus titration by plaque assay both showed similar results (Fig 2I and 2J). Taken together, these data suggested that ASFV p17 could inhibit the cGAS-STING signaling mediated anti-HSV1 and anti-VSV responses.

### 2.3 ASFV p17 is located in ER and Golgi apparatus, and co-localized with STING protein

In order to investigate the molecular mechanism of p17 inhibition of cGAS-STING pathway, the subcellular localization of p17 in cells, and the co-localization between p17 and related proteins of cGAS/STING pathway were assessed by immune fluorescence assay (IFA). The results from IFA showed that p17 protein was co-localized with the ER retention RFP marker, and Golgi apparatus RFP marker, but not with the mitochondria-targeting GFP marker and lysosome LAMP1-GFP marker (SupFig 1A-D). The results also suggested that p17 was co-localized with STING protein, which is a ER resident protein, but no co-localization with cGAS, TBK1, IKKε, IRF3, IFI16 and p65 (Fig 3A-F). Taken together, these dada suggested that p17 is located in ER and Golgi apparatus, and co-localized with STING protein.

**Figure 3.**
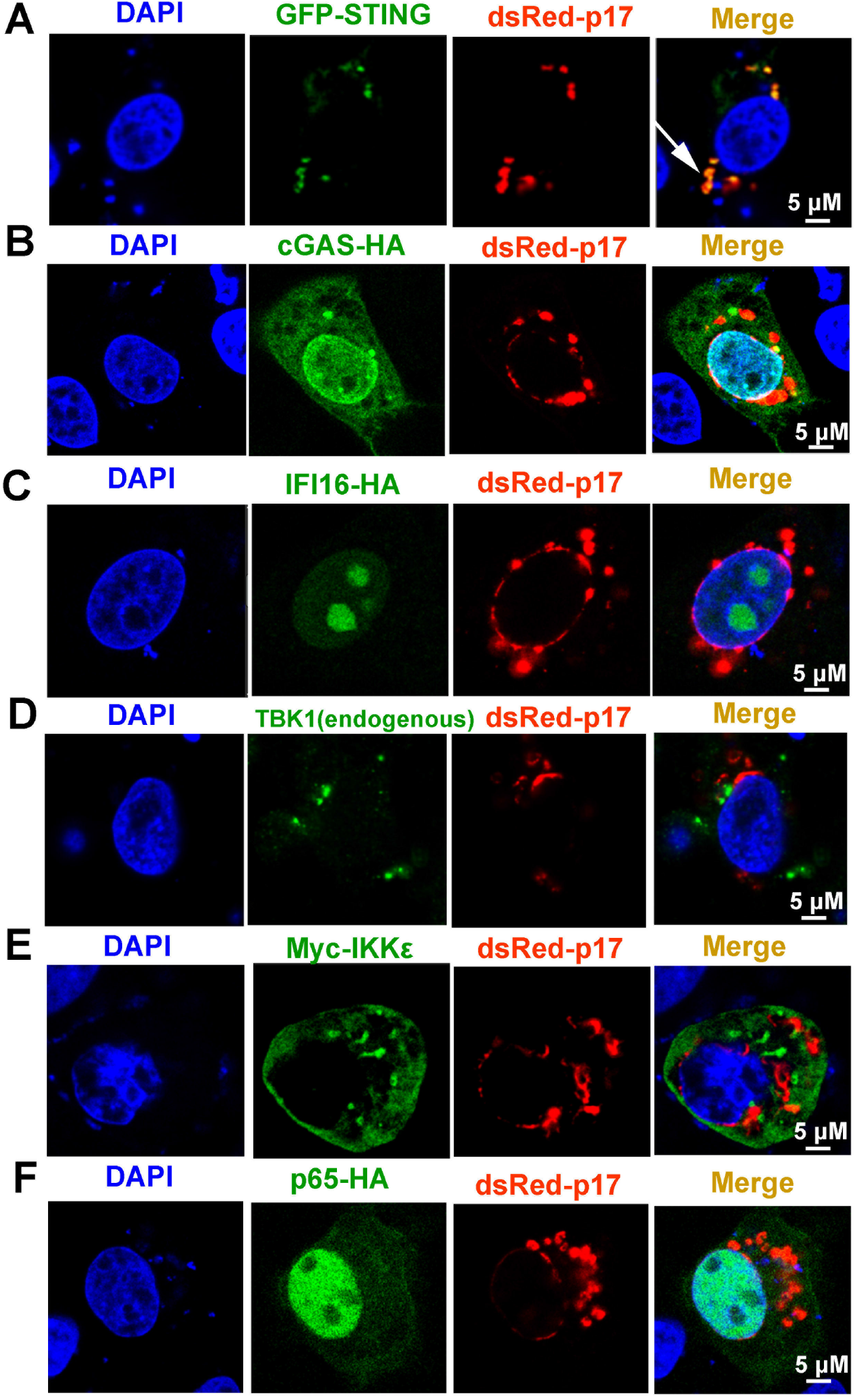
The cellular co-localizations between ASFV p17 and the related proteins of cGAS-STING pathway. PAMs grown on glass coverslip in 24-well plate (1.5×10^5^ cells/well) were co-transfected with dsRed-p17 (0.5 μg) plasmids plus GFP-STING (0.5 μg) (A), cGAS-HA (0.5 μg) (B) IFI16-HA (0.5 μg) (C), Myc-IKKε (0.5 μg) (E) and p65-HA (0.5 μg) (F), respectively, as indicated. HA tagged proteins were stained with rabbit anti-HA antibody and Alexa Fluor plus 488 anti-mouse second antibody. Myc-IKKε was stained with rabbit anti-Myc antibody and Alexa Fluor plus 488 anti-rabbit second antibody. (D) Endogenous TBK1 was stained with rabbit anti-TBK1 antibody and Alexa Fluor plus 488 anti-rabbit second antibody. GFP and dsRed expressions in cells were directly visualized by confocal microscopy. The cellular co-localizations of p17 protein with various cGAS-STING pathway proteins in PAMs was visualized by confocal fluorescence microscopy. The arrow illustrates the co-localized punctate pattern.

### 2.4 p17 inhibits the cGAS-STING pathway through its interaction with STING and interference with STING in the recruitment of TBK1 and IKKε

Considering the cellular co-localization of ASFV p17 and STING, we hypothesized that ASFV p17 likely interferes with STING to affect the cGAS-STING pathway. Thus, we detected the interaction between p17 and STING protein using Co-IP and the results showed that p17 could interact with STING (Fig 4A). Co-IP assay also showed that p17 did not interact with either TBK1 or IKKε protein (Fig 4B-D). To confirm and explore the roles of TBK1 and IKKε in the cGAS-STING signaling, the interactions between STING and these two signaling proteins were also examined by Co-IP. TBK1 was well-defined in mediating STING signaling, and the results showed that upon STING activation, STING was able to interact with endogenous TBK1 as expected (Fig 4E). On the other hand, the role of IKKε in STING signaling has not been well-appreciated, and our results showed that STING also interacted with IKKε (Fig 4F). The data from IFA showed that STING co-localized with TBK1 and IKKε protein as punctate patterns confirming the Co-IP results (Figure 4G and 4H).

**Figure 4.**
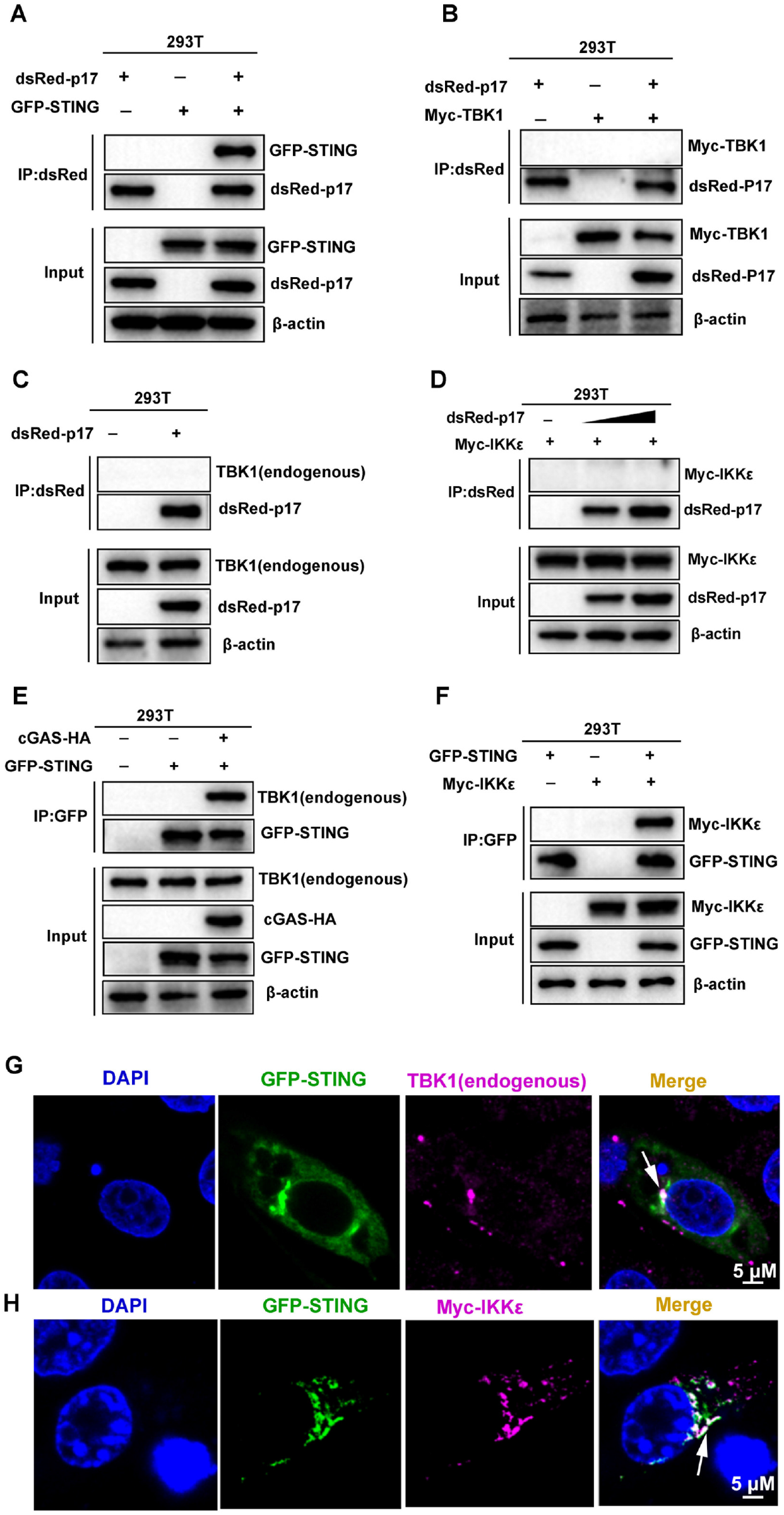
The interactions and cellular co-localizations between p17, STING, TBK1 and IKKε. (A-F) Co-immunoprecipitation assays for analysis of protein interactions. (A) DsRed-p17 (0.5 μg) and GFP-STING (0.5 μg) plasmids were co-transfected into 293T cells in 6 well plate (6-8×10^5^ cells/well) for 24 h. (B) DsRed-p17 (0.5μg) and Myc-TBK1 (0.5 μg) plasmids were co-transfected into 293T cells for 24 h. (C) DsRed-p17 (1 μg) plasmid was transfected into 293T cells for 24 h. (D) DsRed-p17 (0.5 μg and 1 μg) and Myc-IKKε (0.5 μg) plasmids were co-transfected into 293T cells for 24 h. (E) The GFP-STING (0.5μg) and cGAS-HA (0.5 μg) plasmids were transfected into 293T cells for 24 h. (F) The GFP-STING (0.5 μg) and Myc-IKKε (0.5 μg) plasmids were co-transfected into 293T cells for 24 h. The cell lysates were subjected for immuno-precipitation and subsequent Western blotting using the indicated antibodies. (G) The plasmid of GFP-STING (1 μg) were transfected into PAMs for 12h and then stimulated by transfection of polydA:dT (1 g/mL) for another 12h. The endogenous TBK1 was stained by rabbit anti-TBK1 mAb and Alexa Fluor plus 647 anti-rabbit second antibody, and the stained cells were examined by confocal microscopy. (H) The plasmids of GFP-STING (0.5 μg) and Myc-IKKε (0.5 μg) were co-transfected into PAMs for 12 h and then stimulated by transfection of polydA:dT for another 12 h. The Myc-IKKε was stained with mouse anti-Myc pAb and Alexa Fluor plus 647 anti-rabbit second antibody, and stained cells were observed by confocal microscopy. The cellular co-localizations between STING and TBK1/IKKε were indicated by arrows.

Based on above results, we explored the effects of ASFV p17 on the interaction between STING and TBK1/IKKε, Co-IP results showed that p17 was able to inhibit not only the interaction of STING and TBK1 (Fig 5A), but also the interaction of STING and IKKε (Fig 5B). The IFA assay also suggested that in the presence of p17, the co-localizations between STING and TBK1 (Fig 5C), and between STING and IKKε disappeared (Fig 5D).

**Figure 5.**
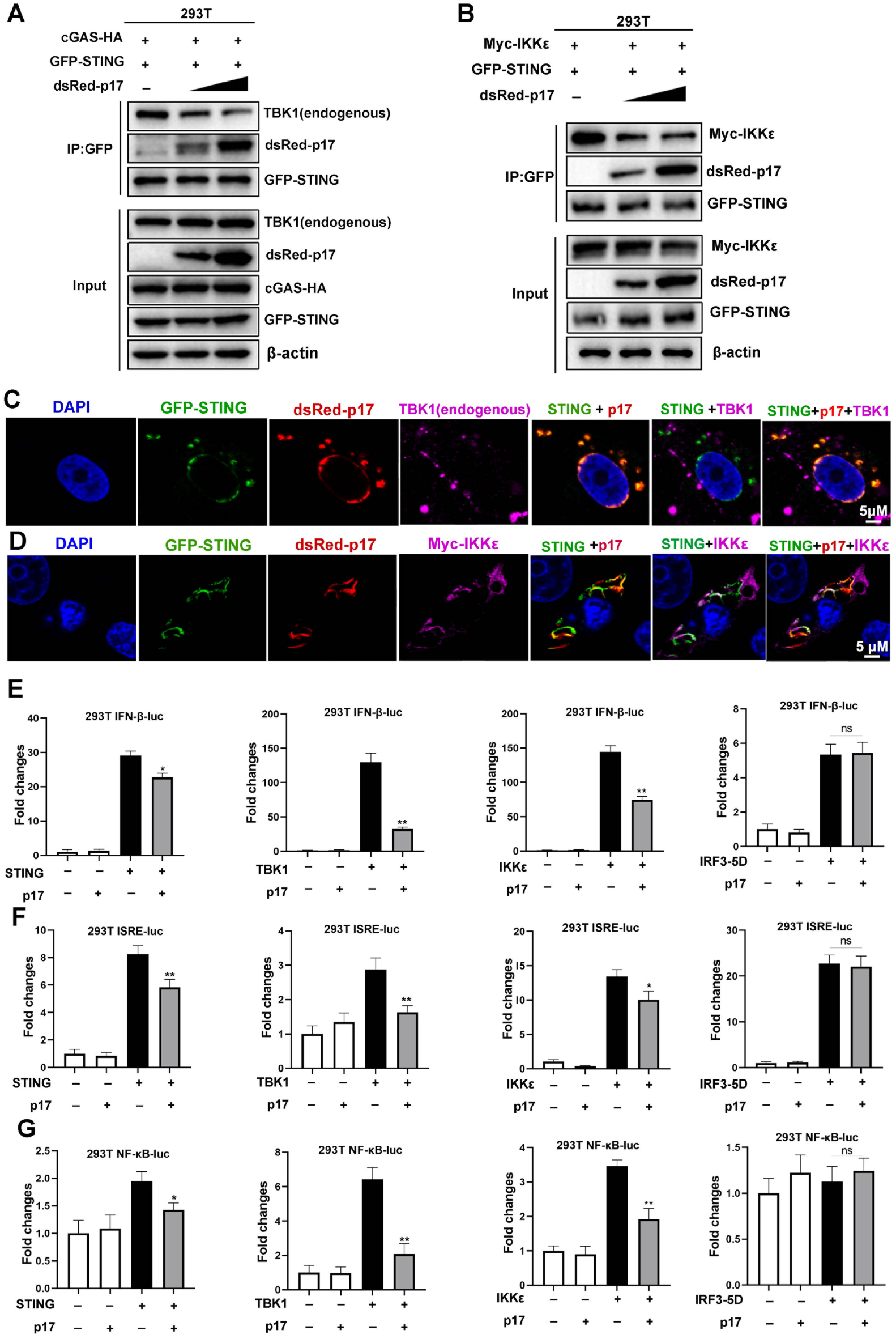
The effects of p17 on the interactions and cellular co-localizations between STING and TBK1/IKKε. (A) 293T cells in 6 well plate (6-8×10^5^ cells/well) were co-transfected with cGAS-HA (0.5 μg) and GFP-STING (0.5 μg) with or without dsRed-p17 (0.5 μg and 1 μg) for 24 h. (B) 293T cells were co-transfected with plasmids of GFP-STING (0.5 μg) and Myc-IKKε (0.5 μg) with or without dsRed-p17 (0.5 μg and 1 μg) for 24 h. The cell lysates were subjected for immuno-precipitation and Western blotting using the indicated antibodies. (C) The effect of p17 on the cellular co-localization between transfected STING and endogenous TBK1 was analyzed by IFA. GFP and dsRed expression in cells was directly visualized by confocal microscopy. The endogenous TBK1 was stained by rabbit anti-TBK1 mAb and Alexa Fluor plus 647 anti-rabbit secondary antibody, and the stained cells were examined by confocal microscopy. (D) The effect of p17 on the cellular co-localization between transfected STING and Myc-IKKε was analyzed by IFA. The Myc-IKKε was stained with rabbit anti-Myc pAb and Alexa Fluor plus 647 anti-rabbit secondary antibody, and stained cells were observed by confocal microscopy. (E-G) The effects of p17 on the activations of IFN promoter (E), ISRE promoter (F) and NF-κB promoter (G) triggered by STING (20 ng), TBK1 (20 ng), IKKε (20 ng) and IRF3-5D (20 ng), which were measured by dual-luciferase reporter assay.

To determine the signaling target of p17 in the inhibition of the cGAS-STING pathway, p17 and the signaling molecules STING, TBK1, IKKε or IRF3-5D were co-transfected into 293T cells and downstream promoter activities examined. The results showed that p17 could inhibit the STING, TBK1 and IKKε but not IRF3-5D mediated IFNβ (Fig 5E), ISRE (Fig 5F) and NF-κB (Fig 5G) promoter activations. This results together with the interaction results suggested that ASFV p17 could inhibit the cGAS-STING pathway through its interactions with STING and interference with STING recruitment of TBK1 and IKKε, thus dampening not only STING signaling but also TBK1/IKKε signaling.

### 2.5 The transmembrane (amino acids 39–59) of p17 is required for its interaction with STING and inhibition of cGAS-STING pathway

The protein sequence analysis of p17 from Uniprot website (https://www.uniprot.org) showed that there are three glycosylation sites including N12, N17 and N97, and one transmembrane domain (A39-Y59) in p17 (Fig 6A). To pinpoint the individual roles of each functional site in the inhibitory activity, three point mutants N12A, N17A and N97A, and one deletion mutant Δ39-59 were made. The results from IFA showed that three mutants N12A, N17A and N97A co-localized with STING, but Δ39-59 had no co-localization with STING (Fig 6B). The promoter assay in 293T cells showed that three mutants N12A, N17A and N97A could inhibit the activation of ISRE promoter (Fig 6C), whereas Δ39-59 had no effect on the activation of IFNβ, ISRE and NF-κB promoters induced by cGAS-STING pathway (Fig 6D-F). Similarly, the results from PAMs have indicated that Δ39-59 had no effects on the activation of IFN-β, ISRE and NF-κB promoters induced by both polydA:dT and 2’3’-cGAMP, respectively (Fig 6G and 6H). The effects of Δ39-59 on the polydA:dT activated cellular gene transcriptions were analyzed by RT-qPCR, and downstream mRNA expressions of IFN-β, ISG15, ISG56 and IL-8 were affected by p17 but not by mutant Δ39-59 (Fig 6I). These data suggested that the transmembrane of p17 is required for the co-localization with STING and the inhibition of cGAS-STING pathway.

**Figure 6.**
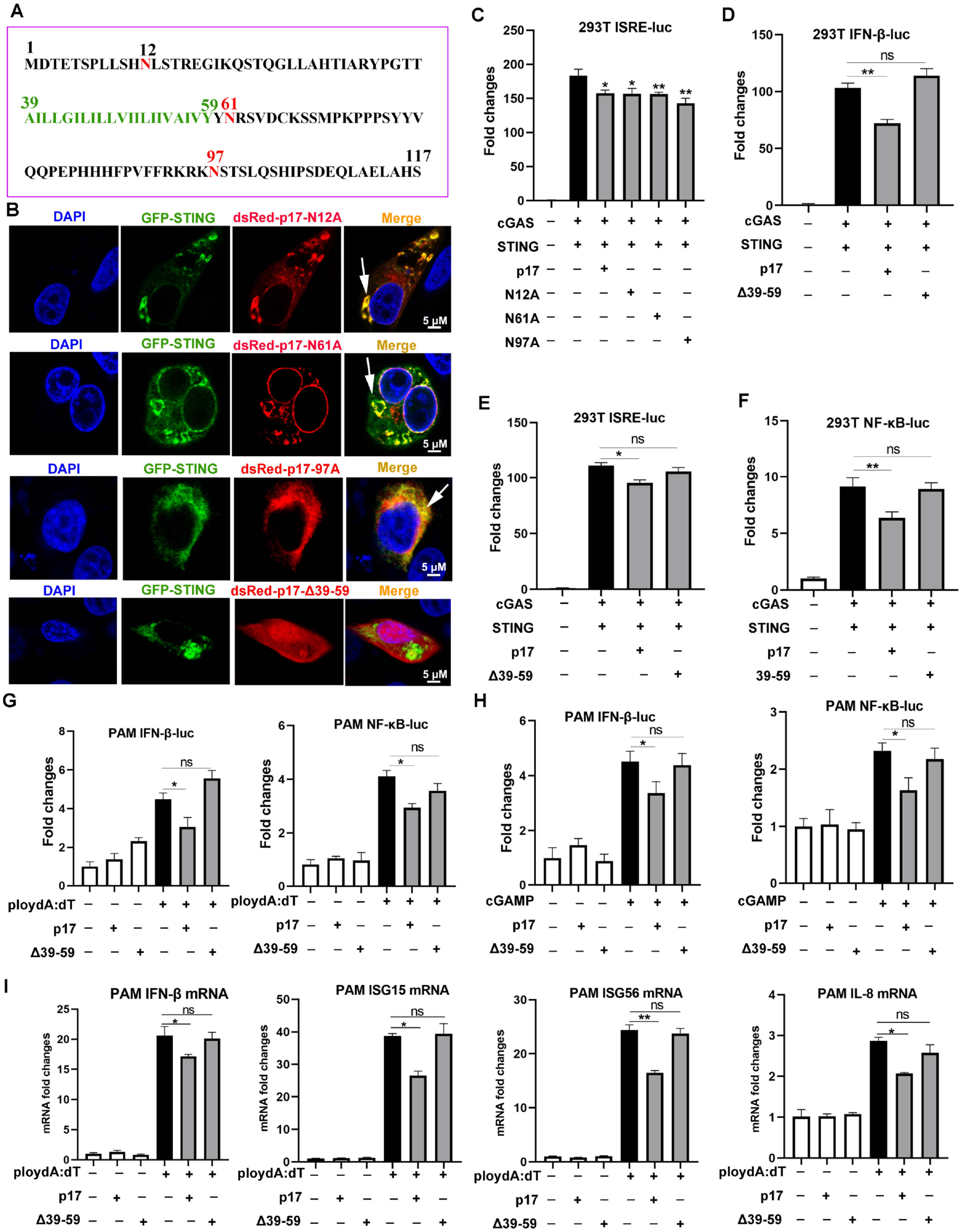
The effects of p17 mutants on cGAS-STING signaling pathway. (A) The protein amino acid sequence of p17, which consists of 117 amino acids and includes three glycosylation sites (red marked N12, N17 and N97) and one transmembrane domain (green marked A39-Y59). (B) The co-localizations between STING and p17 N12A, N17A, N97A, Δ39-59 were examined by confocal microscopy. (C) The effects of p17 N12A, N17A and N97A on the ISRE promoter activation triggered by transfected porcine cGAS-STING in 293T cells was measured by Double-luciferase reporter assay. (D-F) The effects of p17 Δ39-59 on the promoter activations of IFN-β, ISRE and NF-κB triggered by cGAS-STING pathway in transfected 293T cells. The arrows represent the co-localized punctate patterns. (G and H) The effects of Δ39-59 on the activations of promoter IFN-β and NF-κB triggered by transfection of polydA:dT (1 μg/mL) and cGAMP (2 μg/mL) in PAMs. (I) The effect of Δ39-59 on the polydA:dT activated mRNA expressions of IFN-β, ISG15, ISG56 and IL-8 in PAMs assayed RT-qPCR.

We further investigated the impact of mutant Δ39-59 on the interaction of STING and TBK1/IKKε and the signaling triggered by these molecules. The Co-IP results showed that, compared with p17, the Δ39-59 lost the ability to disturb the interaction of STING and TBK1 (Fig 7A), the interaction of STING and IKKε (Fig 7B). Consistently, relative to p17, mutant Δ39-59 also lost the ability to inhibit the signaling activities triggered by STING, TBK1 and IKKε in IFN-β promoter assay (Fig 7C), in ISRE promoter assay (Fig 7D), and in NF-κB promoter assay (Fig 7E).

**Figure 7.**
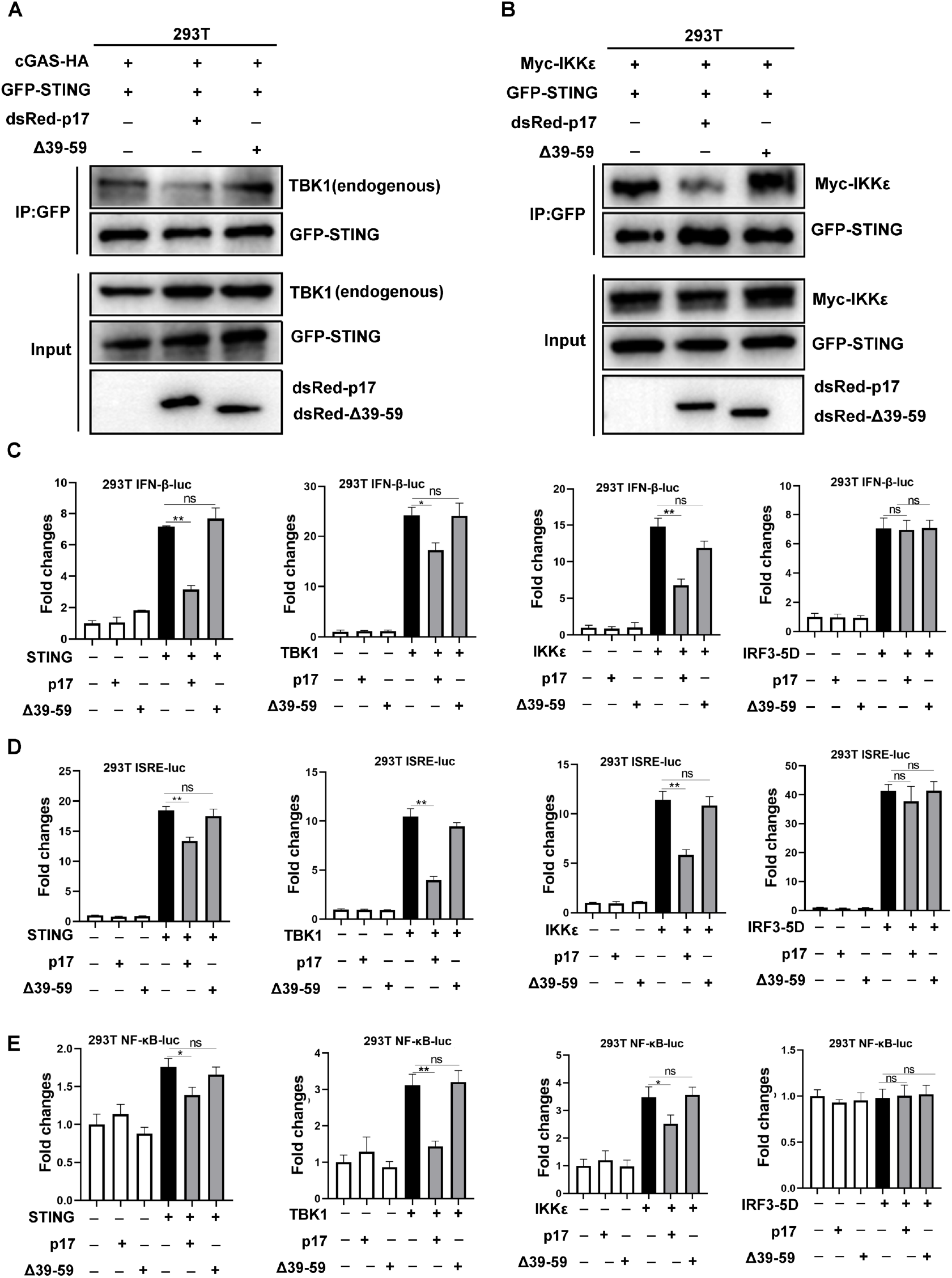
The effect of p17 Δ39-59 on the binding between STING and TBKl/IKKε. (A) 293T cells in 6 well plate (6-8×10^5^ cells/well) were transfected with cGAS-HA (0.5 μg) and GFP-STING (0.5 μg) plus dsRed-p17 (0.5 μg) or Δ39-59 (0.5 μg) for 24h (B) 293T cells were transfected with GFP-STING (0.5 μg) and Myc-IKKε (0.5 μg) plus dsRed-p17 (0.5 μg) or Δ39-59 (0.5 μg) for 24h. The effect of p17 Δ39-59 on the binding between STING and TBK1/IKKε was analyzed by Co-IP and Western blotting using the indicated antibodies. (C-E) The effects of p17 Δ39-59 on the activations of IFN-β (C), ISRE (D) and NF-κB (E) promoter triggered by STING (20 ng), TBK1 (20 ng), IKKε (20 ng) and IRF3-5D (20 ng) were examined using dual-luciferase reporter assay.

Additionally, we found that Δ39-59 lost the ability to inhibit the 2’3’-cGAMP-STING signaling mediated anti-HSV1 activity in PAMs, as evidenced by fluorescence microscopy (Fig 8A), flow cytometry (Fig 8B) and Western blotting (Fig 8C). Similarly, in 293T cells, Δ39-59 lost the ability to inhibit the porcine cGAS-STING signaling mediated anti-VSV activity, as evidenced by fluorescence microscopy (Fig 8D), flow cytometry (Fig 8E) and Western blotting (Fig 8F). Taken together, all these data suggested that the transmembrane of p17 is required for its STING interaction, disruption of the interaction between STING and TBK1/IKKε, and thus suppression of the triggered signaling activates and antiviral responses.

**Figure 8.**
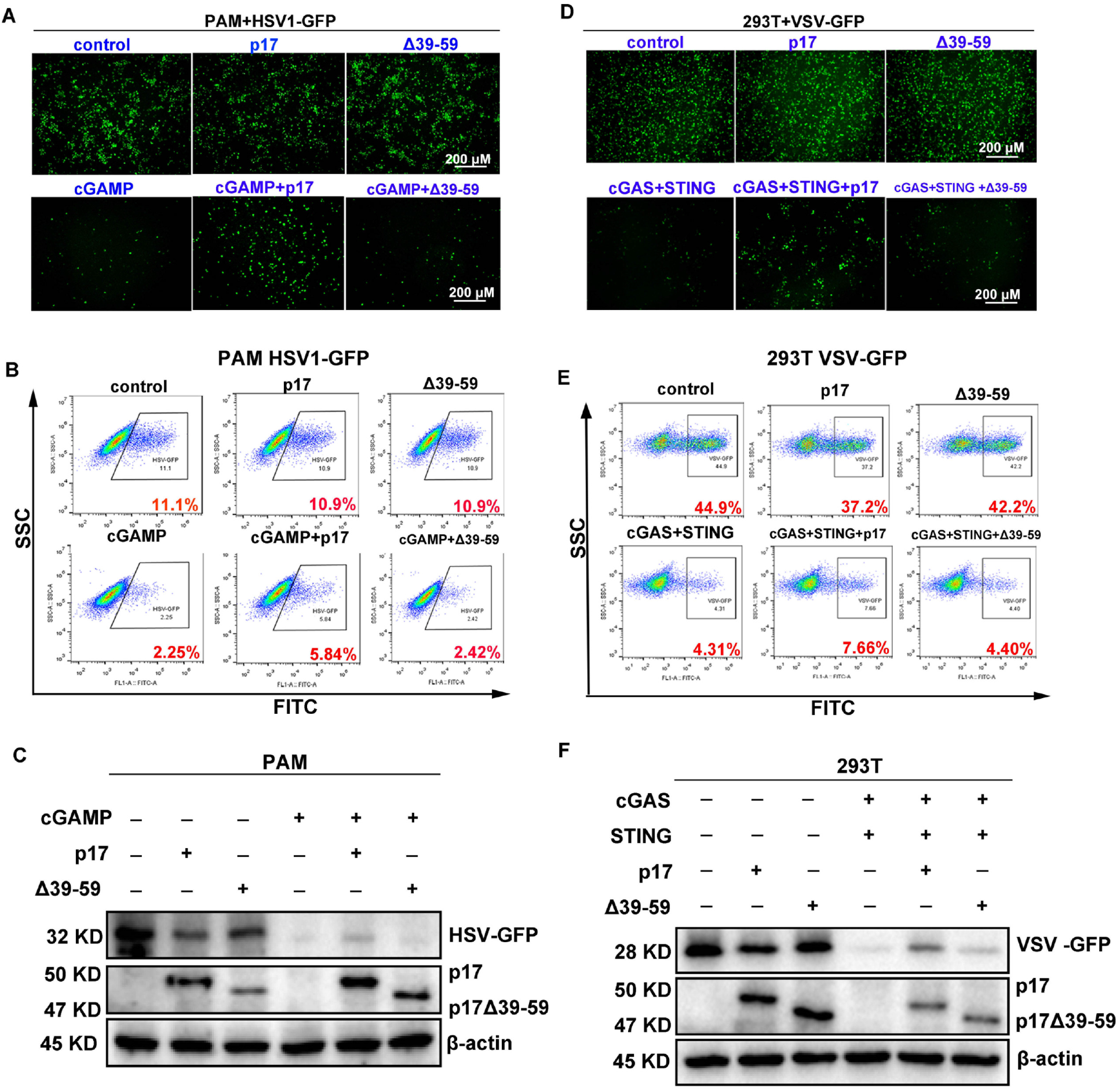
Effects of p17 Δ39-59 on the cGAS-STING signaling mediated anti-HSV1 and anti-VSV responses. (A) The p17 (0.5 μg) or Δ39-59 (0.5 μg) was transfected into PAMs for 12 h, and the stimulated with transfection with 2’3’-cGAMP (2 μg/mL) for 12h. The cells were infected with HSV1-GFP at the MOI of 0.01 for 20 h. (A) The HSV1 replicative GFP signals were visualized by fluorescence microscopy. (B) The HSV1-GFP infected cells were analyzed by flow cytometry. (C) The viral GFP protein expressions were detected by Western blotting. (D-F) The p17 (0.5 μg) or Δ39-59 (0.5 μg) plus porcine cGAS (0.5 μg) and STING (0.5μg) were co-transfected into 293T cells for 12 h. The cells were infected with VSV-GFP at the MOI of 0.001 for 8 h. The VSV replicative GFP signals were visualized by fluorescence microscopy (C). The infected cells were analyzed by flow cytometry (E). The viral GFP protein expressions were analyzed by Western blotting (F).

### 2.6 Analyzing the amino acid sequences responsible for the co-localization between ASFV p17 and STING

In order to minimize the functional sequence in the transmembrane (AAs 39–59) of p17, more deletion mutants of p17 in pdsRed-Express-C1, including Δ39-48, Δ44-53, Δ49-59, Δ39-43, Δ44-48, Δ49-53 and Δ53-59 were made. The results from IFA showed that Δ39-48, Δ44-53, Δ49-59 and Δ53-59 had no co-localization with STING protein as Δ39-59 (SupFig 2A). However, Δ39-43, Δ44-48, Δ49-53 were still co-localized with STING protein (SupFig 2A), indicating that most sequences except amino acids 39-43 of p17 is indispensable for its interaction with STING.

On the other hand, the roles of each functional domains of porcine STING in the interaction with ASFV p17 were also examined by IFA. The porcine STING protein includes four functional domains which are N-terminal domain mediating interaction with ZDHHC1 and ZDHHC11 (AAs 1-190), cyclic dinucleotide-binding domain (AAs 153-339), c-di-GMP binding domain (AAs 238-241) and C-terminal tail (AAs 339-378) (SupFig 2B). Four deletion pSTING mutants Δ1-190, Δ153-339, Δ238-241 and Δ339-378, and three individual pSTING fragments 1-190, 153-339 and 339-378 were obtained as pEGFP-C1 recombinant vectors, respectively. The con-focal microscopy results showed that Δ238-241 and Δ339-378 were co-localized with ASFV p17, however, Δ1-190, Δ153-339 and all three individual fragments lost the co-localization with ASFV p17. These data indicated the N-terminal domain (AAs 1-190) and middle domain (AAs 153-339) of STING are both required for the co-localization with ASFV p17 and STING (SupFig 2C).

## 3. Discussion

Viral infection triggers a series of signaling cascades that results in the expression of type I IFNs, which exerts key roles in cellular antiviral responses (32, 33). The cGAS-STING DNA sensing pathway has been reported to play a pivotal role in inducing the production of IFNs and suppressing the replication of viruses, in particular DNA viruses (34–36). However, viruses have developed multiple strategies to resist host immune defenses, and counteract IFN signaling (37). Recently, studies have indicated that Armenia/07 virulent strain mediated production of IFN-β through the cGAS-STING pathway (15), whereas the molecular mechanism of ASFV interacting with the cGAS-STING pathway has not been fully elucidated. In the current study, our results demonstrated that ASFV structural protein p17 exerts a negative regulatory effect on cGAS-STING signaling pathway and the signaling dependent antiviral responses by targeting STING. To our knowledge, this is the first time that ASFV p17 is reported to inhibit cGAS-STING pathway.

ASFV p17 protein is encoded by the D117L gene and consists of 117 amino acids. There are three glycosylation sites including N12, N61 and N97, and one transmembrane domain (AAs 39-59) (23). We found that the transmembrane domain of p17 is required for interaction with STING and interference with STING in the recruitment of TBK1 and IKKε while the three glycosylation sites are not important. The transmembrane domain of p17 is also critical for the subversion of cGAS-STING signaling mediated anti-HSV1 and anti-VSV responses. Intriguingly, p17 alone during HSV1 and VSV infections did not promote the viral replications as expected, instead, it weakly inhibited the viral replications, suggesting p17 is a multiple functional protein. Indeed, ASFV p17 was able to suppress cell proliferation via ER-ROS pathway as shown by us recently (23), and p17 is likely to suppress cell autophagy (unpublished data). All these p17 cellular functions influence virus replication, and collectively, the inhibition and not the enhancement of virus replications was observed when p17 present alone during virus infection. The three glycosylation sites and transmembrane domain of p17 may play potential roles in the p17 various cellular functions; therefore, what roles of these sites of p17 in its various cellular functions will be interesting and warranted to be further investigated.

Accumulating evidences suggest that STING can be manipulated by several viral products to counteract the cGAS-STING pathway and type I IFN production (38). STING, an ER-associated protein, is essential for inducing IFN in response to intracellular DNA or DNA pathogens including bacteria and DNA viruses (39, 40). In the presence of cytosolic DNA which binds with cGAS to produce 2’3’-cGAMP, STING dimer self oliogmerizes, and translocates sequentially from the ER to ERGIC, Golgi apparatus, and eventually relocates to lysosome for degradation (41, 42). Upon translocation from ERGIC on, STING recruits TBK1 to induce IFNs and other cytokines (43). Our study found that ASFV p17 targets STING, disturbs the interaction of STING and TBK1, and thus inhibits IFN response. Not only that, we also found that ASFV p17 also disturbs the interaction of STING and IKKε. One recent study reported that IKKε and TBK1 act redundantly to activate STING mediated downstream NF-κB signaling (44). How significant the IKKε is in STING mediated IFN response and how significant the suppression of IKKε-STING interaction by p17 is in the ASFV immune evasion are not known and need to be addressed in the future.

In addition, our co-localization results indicated that the most transmembrane region of p17 except amino acids 39-43 may be important for interaction with STING. On the other hand, in STING, the N-terminal portion and middle cyclic dinucleotide-binding domain might be required for interaction with ASFV p17, and the C-terminal CTT domain of STING is not involved in the interaction. The CTT domain is the key portion for STING to recruit and activate TBK1 and IRF3, thus the alteration of STING and TBK1 interaction by p17 is likely occurring through an indirect way. The dissection of molecular mechanism controlling p17 and STING interactions provided deep insights into the understanding of ASFV p17 action in immune evasion events. However, the interaction detail needs to be further confirmed by other experiments.

In summary, this study revealed that ASFV p17 is able to inhibit the cGAS-STING pathway through interaction with STING and interference with STING in the recruitment of TBK1 and IKKε (SupFig 3). The transmembrane domain of p17 is required in the interaction with STING and the inhibition of cGAS-STING pathway. Findings in this study will expand our knowledge on the molecular mechanisms by which ASFV counteracts the antiviral innate immunity and provide deep insights into ASF pathogenesis.

## 4. Material and methods

### 4.1. Antibodies and reagents

The rabbit TBK1 mAb (3504S), phosphorylated-TBK1 mAb (p-TBK1, 5483S), IRF3 mAb (11904S), FLAG mAb (14793), HA mAb (3724) and GFP mAb (2956) were acquired from Cell Signaling Technology (Boston, MA, USA). The rabbit p-IRF3 (Ser385) pAb (MA5-14947) was purchased from Thermo Fisher (Sunnyvale, CA, USA). The rabbit STING pAb (19851-1-AP), and MYC pAb (16286-1-AP) were purchased from ProteinTech (Wuhan, Hubei, China). The dsRed mAb (ab185921) was acquired from Abcam (Cambridge, Cambridgeshire, UK). The mouse cGAS mAb (sc-515777) was acquired from Santa Cruz Biotechnology (Santa Cruz, CA, USA). HRP goat anti-rabbit IgG (H+L) highly cross-adsorbed secondary antibody and goat anti-mouse IgG (H+L) highly cross-adsorbed secondary antibody, Goat anti-rabbit IgG (H+L) highly cross-adsorbed secondary antibody, Alexa Fluor Plus 488 (A32731), Alexa Fluor Plus 647 (A32733), and Goat anti-mouse IgG (H+L) highly cross-adsorbed secondary antibody, Alexa Fluor 488 (A11029) were all acquired from Thermo Fisher (Sunnyvale, CA, USA).

DMEM medium and RPMI 1640 medium were obtained from Hyclone (Hyclone Laboratories, Logan, Utah, USA). Fetal bovine serum (FBS) was obtained from Gibco (Grand Island, NY, USA). Double-luciferase reporter assay Kit were bought from TransGen Biotech (Beijing, China). 2’3’-cGAMP and polydA:dT were bought from InvivoGen (Hong Kong, China). Lipofectamine 2000 was bought from Invitrogen (Carlsbad, CA, USA). Protein A/G PLUS-Agarose was bought from Santa Cruz Biotechnology (sc-2003, CA, USA). DAPI staining solution (C1005), Enhanced BCA protein assay kit (P0010S), Cell lysis buffer (P0013) were purchased from Beyotime (Shanghai, China). All other chemicals and reagents were analytical grade and were obtained commercially.

### 4.2. Cells, cell transfection and viruses

Human embryonic kidneys (293T) was maintained in a DMEM medium supplemented with 10% fetal bovine serum, 1 mM glutamine, and 1% penicillin/streptomycin, and maintained at 37 °C with 5% CO_2_. Porcine alveolar macrophages (PAMs, 3D4/21) were maintained in an RPMI 1640 medium supplemented with 10% fetal bovine serum, 1 mM glutamine, and 1% penicillin/streptomycin, and maintained at 37 °C with 5% CO_2_. Transfection was performed by using the Lipofectamine 2000 following the manufacturer’s instructions. The Vesicular Stomatitis Virus (VSV-GFP) and Herpes Simplex Virus-1 (HSV-1-GFP) were both provided by Dr. Tony Wang in SRI International USA, and used as we described previously (31).

### 4.3. Plasmids and molecular cloning

The D117L (p17) gene of ASFV China 2018/1 (GenBank submission No. MH766894) was codon optimized for protein expression in mammalian cells. It was synthesized and cloned into 3xFLAG-CMV-7.1 vector using *Hind* III and *Kpn* I sites, pdsRed-Express-C1 vector using *Bgl* II and *EcoR* I sites, respectively. Three mutants of p17, including N12A, N17A and N97A, and one deletion mutant Δ39-59 were made by mutation PCR with PrimerSTAR Max DNA Polymerase (Takara, Beijing, China) and pdsRed-p17 as the template. Eight deletion mutants of p17 including Δ39-59, Δ39-48, Δ44-53, Δ49-59, Δ39-43, Δ44-48, Δ49-53 and Δ54-59 were made by mutation PCR using pdsRed-p17 as the template.

Porcine STING (pSTING) open reading frame was cloned in *Bgl* II and *EcoR* I sites of pEGFP-C1 and pmCherry-C1 vector, respectively. The pSTING Δ238-241 was obtained by mutation PCR using PrimerSTAR Max DNA Polymerase and pEGFP-pSTING as the template. All the mutation PCR products were digested with *Dpn* I and transformed into competent DMT *E.coli* to obtain the mutant plasmids. For pSTING Δ1-190, Δ339-378, 1-190, 153-339 and 339-378, the fragments were directly amplified from pEGFP-pSTING by PCR. For pSTING Δ153-339, the two fragments flanking the deletion site were amplified by PCR from template pEGFP-pSTING, and next the two flanking fragments were joined together by the fusion PCR. The PCR products were then cloned into *Bgl* II and *EcoR* I sites of pEGFP-C1, respectively, as the template pEGFP-pSTING.

The mutation PCR primers of p17 and pSTING were designed using QuickChange Primer Design method (https://www.agilent.com), which are available with other primers in Sup Table 2. Porcine cGAS, IFI16, IKKβ and p65 were cloned and used as we described before (45). Human TBK1 and IKKε were cloned and preserved in our lab.

### 4.4. Promoter-driven luciferase reporter gene assay

293T cells grown in 96-well plates (3×10^4^ cells/well) were co-transfected by Lipofectamine 2000 with IFB-β-luciferase, ISRE-luciferase or NF-κB-luciferase reporter (10 ng/well) (Fluc) and β-actin *Renilla* luciferase (Rluc) reporter (0.2 ng/well), together with the indicated plasmids or vector control (5–40 ng/well). The total DNA per well was normalized to 50 ng by adding empty vector. After 24 h post-transfection, cells were harvested and lysed using lysis buffer for 15 min at room temperature (RT). Relative luciferase activity was measured using a Double-luciferase reporter assay Kit following the manufacturer’s suggestions. The relative luciferase activity was analyzed by normalizing Fluc to Rluc activity. The results were expressed as fold induction of IFN-β-Fluc, ISRE-Fluc or NF-κB-Fluc compared with vector control group after Fluc normalization by corresponding Rluc.

### 4.5. RNA Extraction and RT-qPCR

Total RNA was extracted by using TRIpure reagent following the manufacturer’s suggestions. The extracted RNA was reverse transcribed into cDNA using HiScript 1^st^ strand cDNA synthesis kit (Vazyme, Nanjing, China), and then the target gene expressions were measured using quantitative PCR with SYBR qPCR master Mix (vazyme, Nanjing, China) in StepOnePlus equipment (Applied Biosystems). The qPCR program is denaturation at 94°C for 30 s followed by 40 cycles of 94°C for 5 s and 60°C for 30 s. The relative mRNA levels were normalized to β-actin mRNA levels, and calculated using 2^−ΔΔCT^ method. The sequence of primers used are shown in Supplementary Table 1.

### 4.6. Western blotting and Co-immunoprecipitation analysis

Whole cell proteins were extracted with an RIPA lysis buffer. Then, the concentration of the whole protein was analyzed and adjusted using the BCA protein assay kit (Beyotime Institute of Biotechnology, Shanghai, China). The protein samples were mixed with 1×loading buffer and boiled for 10 min. The protein supernatants were run by SDS-PAGE, and then the proteins in gel were transferred to PVDF membranes. The membranes were incubated with 5% skim milk solution at RT for 2 h, probed with the indicated primary antibodies at 4°C overnight, washed, and then incubated with secondary antibodies for 1 h at RT. The protein signals were detected by ECL detection substrate and imaging system.

For co-immunoprecipitation (Co-IP), the cleared cell lysate from transfected cells in 6 well plate (6-8×10^5^ cells/well) was incubated with 1 μg/mL of specific antibody at 4°C overnight with shaking and further incubated with Protein A/G PLUS-Agarose for 2-3 h. The agarose was washed and eluted with 40 μL of 2 × SDS sample buffer. The elution samples together with input controls were both subjected to Western blotting.

### 4.7. Confocal microscopy

Cells cultured in cell coverslip in a 24-well plate (2×10^5^ cells/well) were fixed in 4% paraformaldehyde for 30 min at RT, permeabilized by 0.5% Triton X-100, and then blocked with 5% BSA. The treated cells were incubated with primary antibody against FLAG, HA or TBK1 (1: 500) overnight, and then incubated with secondary antibody (1: 500) for 1 h. Finally, the coverslips were counterstained with DAPI, loaded on slide, sealed by nail polish, and visualized under a confocal laser-scanning microscope (Leica TCS SP8,Leica, Weztlar, Germany). Green fluorescence protein GFP and red fluorescence protein dsRed expression in cells was directly visualized by confocal microscopy.

### 4.8. Flow cytometry

PAMs seeded in 12-well plates (3×10^5^ cells/well) were transfected with p17 plasmid (1 μg) for 24 h, and next stimulated by transfection with 2’3’-cGAMP (1 μg/mL) for 12 h. The cells were then infected with HSV1-GFP (MOI 0.01) for 20 h. 293T cells seeded in 12-well plates (3×10^5^ cells/well) were transfected with p17 (0.5 μg) together with cGAS (0.5 μg) and STING (0.5 μg) plasmids for 12 h, and then infected with VSV-GFP (MOI 0.001) for 8 h. After infection, cells were harvested and washed twice with PBS. The cell samples were filtered using a 200-mesh nylon filter, and cell suspensions analyzed by flow cytometry at a wavelength pair of 488/525 nm.

### 4.9. Plaque assay

Vero cells were seeded into 12-well plates (3×10^5^ cells/well). After the cells were grown into monolayer, cells were infected by the tenfold serially diluted cell supernatants from HSV1 or VSV infected cells for 2 h. Then the infected cells were washed with PBS and overlaid by immobilizing medium of 1:1 mixture of warmed 2×DMEM with 4% FBS and a stock solution of heated 1.6% low melting agarose. Plaque formation required 4 days for HSV1, and 2 days for VSV, respectively. Upon completion, the immobilizing medium were discarded by tipping and cells were fixed and stained with crystal violet cell colony staining solution (0.05% w/v crystal violet, 1% formaldehyde, 1×PBS and 1% methanol) for 1 h at room temperature. After staining, cells were washed with tap water until the clear plaques appeared. The plaques were counted and photos were taken.

### 4.10. Statistical analysis

All of the experiments were representative of two or three similar experiments. The results were analyzed using SPSS and presented as the mean ± standard deviation (SD). Statistical analysis was performed using Student’s *t*-test and *ANOVA*; *p* < 0.05 was considered statistically significant. In the figures, “*”, “**” and “ns” denote *p* < 0.05, *p* < 0.01 and statistically not significant, respectively.

---

**Supplementary Figure 1.**
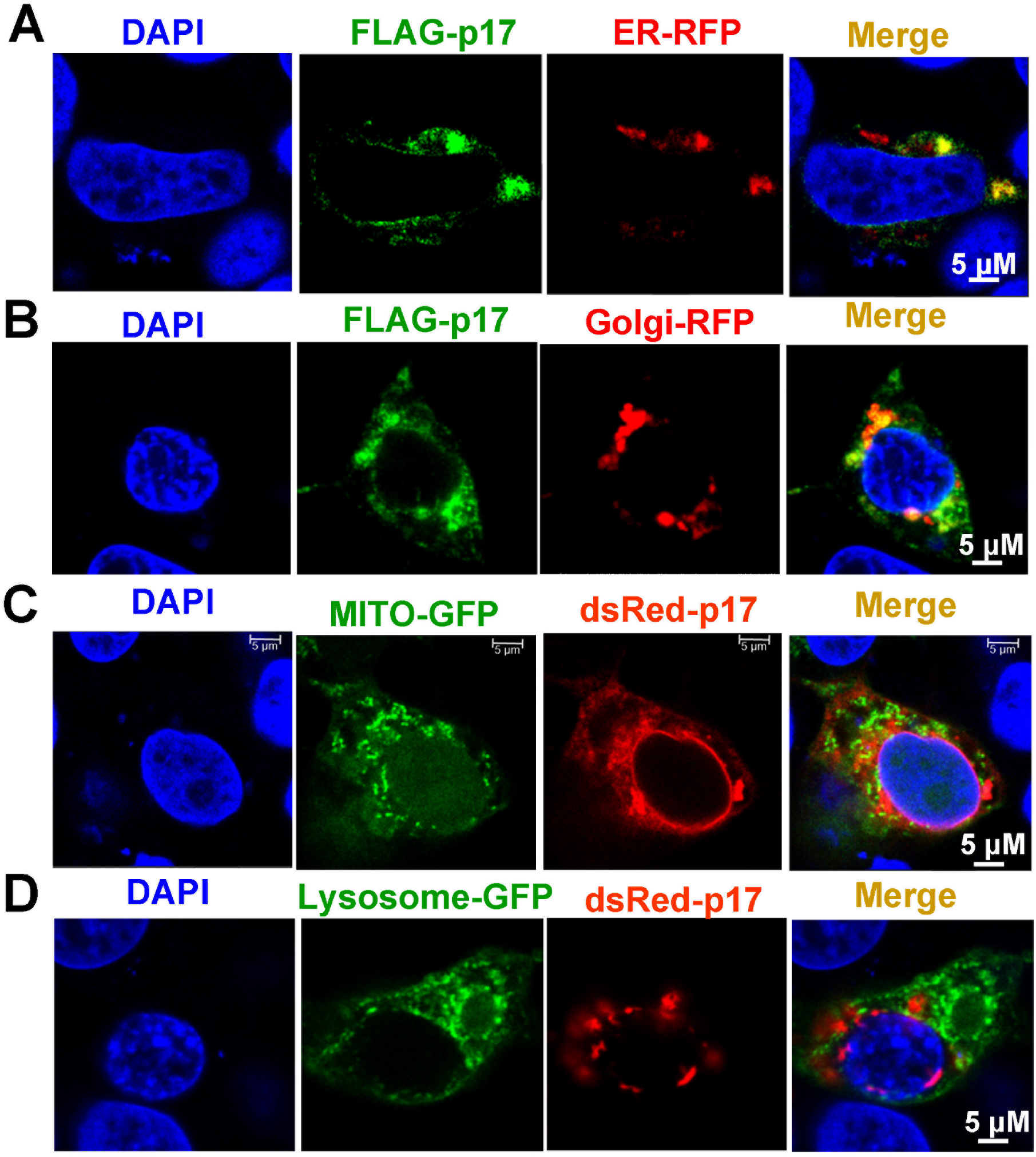
The subcellular localization of p17 in PAMs. PAM cells (1.5×10^5^ cells/well) grown on glass coverslip in 24-well plate were transfected with p17 plasmids (0.5 μg) together with RFP-KDEL (an ER retention protein) (0.5 μg) (A), with Golgi-RFP (0.5 μg) (B), with Mito-EGFP (mitochondrial targeting sequence of cytochrome c oxidase subunit VII fused to EGFP) (0.5 μg) (C), with GFP-LAMP1 (lysosomal-associated membrane protein1) (0.5 μg) (D), respectively, as indicated. The FLAG-p17 was stained with rabbit anti-FLAG mAb and Alexa Fluor plus 488 anti-rabbit second antibody. The subcellular localization of p17 protein in PAMs was visualized by confocal fluorescence microscopy.

**Supplementary Figure 2.**
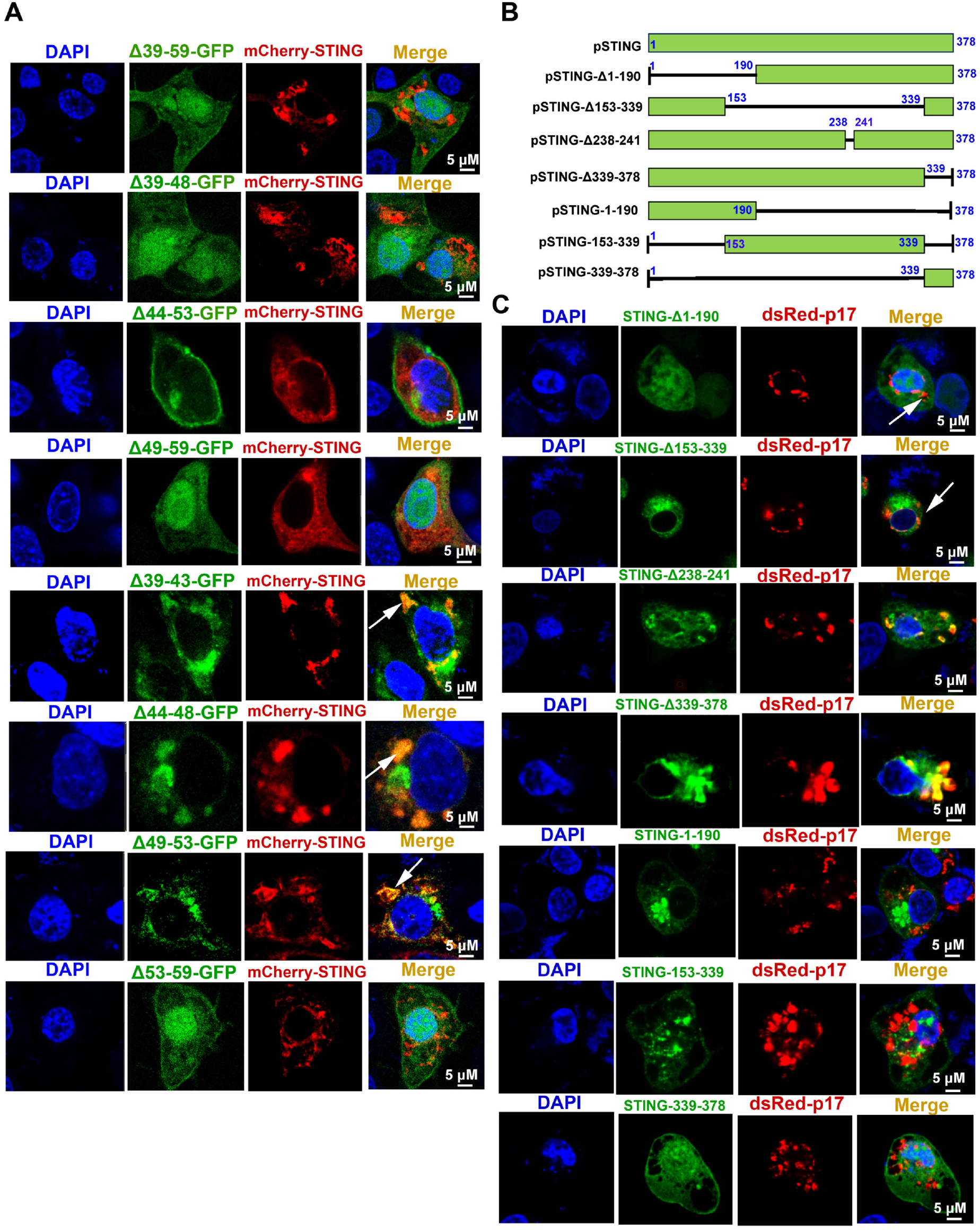
Analyzing the amino acid sequence responsible for the co-localization between p17 and STING. (A) PAMs on coverslip in 12-well plate (1.5×10^5^ cells/well) were co-transfected with mCherry-STING (0.5 μg) plus p17-GFP Δ39-59, Δ39-48, Δ44-53, Δ49-59, Δ39-43, Δ44-48, Δ49-53 or Δ53-59 (0.5 μg each) for 24 h. Cells were fixed, and then counter stained by DAPI. (B) Based on the protein function domains of porcine STING, a series of STING deletion mutants were constructed as indicated. (C) PAMs on coverslip in 12-well plate (1.5×10^5^ cells/well) were co-transfected with dsRed-p17 (0.5 μg) plus GFP-pSTING Δ1-190, Δ153-339, Δ238-241, Δ339-378, 1-190, 153-339 or 339-378 (0.5 μg each) for 24 h, cells were fixed, and then counter stained by DAPI. Green fluorescence protein GFP and red fluorescence protein dsRed expression in PAMs were directly visualized and cellular co-localizations detected by confocal microscopy.

**Supplementary Figure 3.**
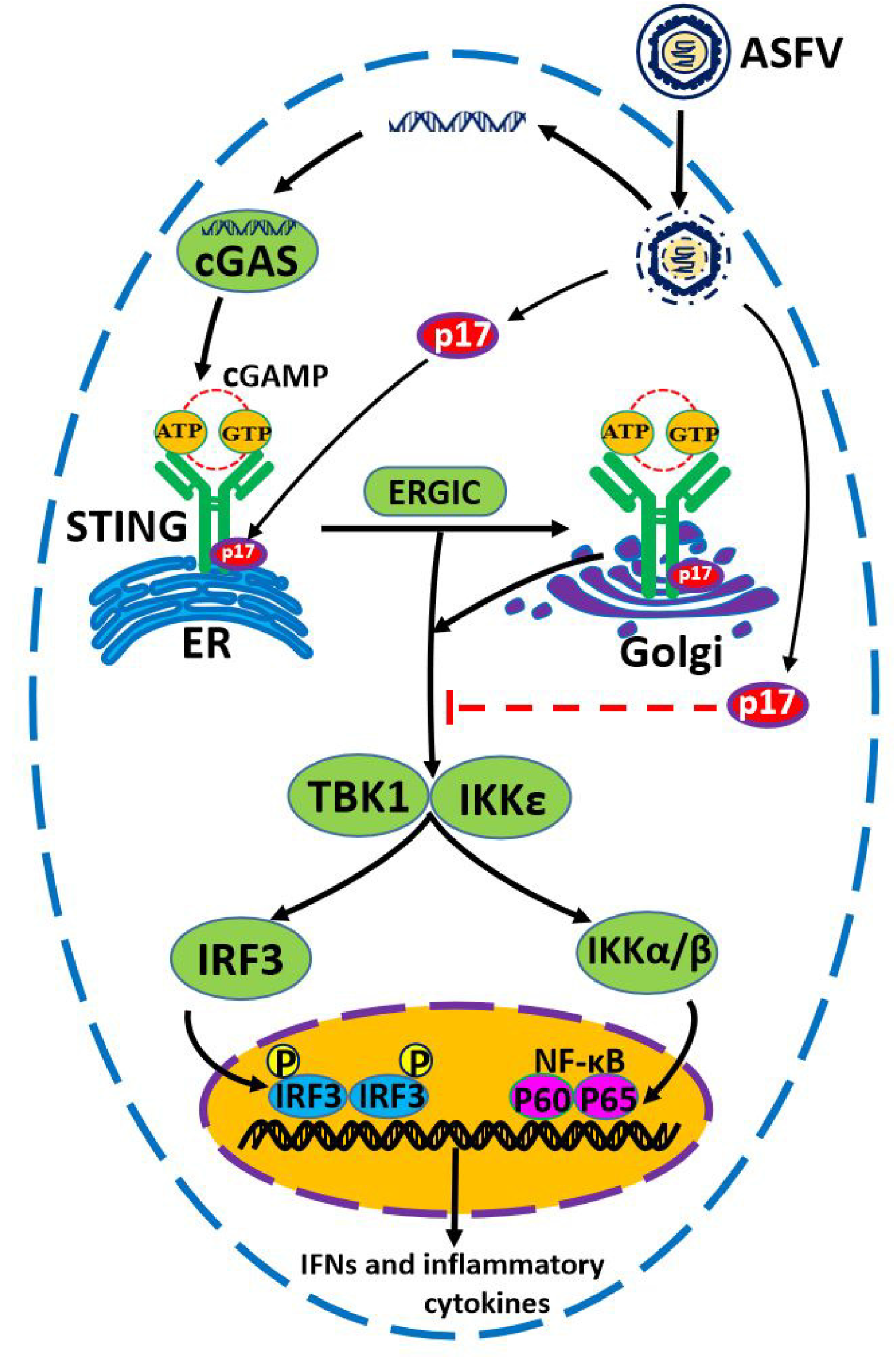
The proposed molecular mechanism of p17 inhibiting innate immune DNA sensing cGAS-STING pathway. Note, ER denotes endoplasmic reticulum and ERGIC denotes ER Golgi intermediate compartment, respectively.

## Author contribution statement

J.Z conceived and designed the experiments; W.Z, N.X, J.L, S.J, J.Z, H.W, D.A, Y.X, Q.S, Q.C, Y.Z performed the experiments; W.Z, J.D, N.C, Q.Z, H.C, X.G, H.Z, F.M, J.Z analyzed the data; W.Z and J.Z wrote the paper. All authors contributed to the article and approved the submitted version.

## Acknowledgments

The work was partly supported by the National Key Research and Development Program of China (2017YFD0502301), Jiangsu provincial key R & D plan (BE2020398), the National Natural Science Foundation of China (31872450), and A Project Funded by the Priority Academic Program Development of Jiangsu Higher Education Institutions (PAPD).

## Data Availability Statement

The data presented in this study are available in insert article.

